# Quantifying the severity of adverse drug reactions using social media

**DOI:** 10.1101/2021.02.02.429445

**Authors:** Adam Lavertu, Tymor Hamamsy, Russ B Altman

## Abstract

Adverse drug reactions (ADRs) impact the health of 100,000s of individuals annually in the United States with associated costs in the hundreds of billions. The monitoring and analysis of the severity of adverse drug reactions is limited by the current qualitative and categorical system of severity classifications. Previous efforts have generated quantitative estimates for a subset of ADRs, but were limited in scope due to the time and costs associated with the efforts. We present a semi-supervised approach that estimates ADR severity by using a lexical network of ADR word embeddings and label propagation. We use this method to estimate the severity of 28,113 ADRs, representing 12,198 unique ADR concepts from MedDRA. Our Severity of Adverse Events Derived from Reddit (Saedr) scores have good correlations with real-world outcomes. Saedr scores had Spearman correlations with ADR case outcomes in FAERS of 0.595, 0.633, and −0.748 for death, serious outcome, and no outcome, respectively. We investigate different methods for defining initial seed term sets and evaluate their impact on severity estimates. We analyzed severity distributions for ADRs based on their appearance in Boxed Warning drug label sections, as well as ADRs with sex-specific associations. We find that ADRs discovered postmarket have significantly greater severity compared to those discovered in the clinical trial. We create quantitative Drug RIsk Profile (Drip) scores for 968 drugs that have a Spearman correlation of 0.377 with drugs ranked by FAERS cases resulting in death, where the given drug was the primary suspect. We make the Saedr and Drip scores publicly available in order to enable more quantitative analysis of pharmacovigilance data.

## Introduction

Adverse drug reactions (ADRs) are one of the leading causes of mortality and morbidity in the United States, affecting hundreds of thousands of people and costing more than $500 billion every year in the United States alone[1,2]. An ADR is characterized as “an appreciably harmful or unpleasant reaction resulting from an intervention related to the use of a medicinal product”[3]. A drug’s ADRs are primarily derived from clinical trial data and augmented through postmarket surveillance[4]. The drug ADR labeling process is based on the frequency of the ADRs in the treated populations and the severity of the outcomes associated with each ADR. The severity of an ADR is traditionally classified into one of three categories characterized by where they appear on the label: Box Warning (BW), Warnings and Precautions (WP), or Adverse Reactions (AR), listed in decreasing order of associated severity[5]. BWs refer to “serious warnings, particularly those that lead to death or serious injury.” Under the existing system, death is severe enough to warrant a BW, but so is “loss of bone density”, while both are serious ADRs, it would be useful for prescribers, patients and researchers to have systems for recognizing that death is more severe than loss of bone density[6]. Similarly, WP and AR include a diverse spectrum of ADRs, and it can be difficult for patients, prescribers and researchers to compare the risk profiles of different drugs. Pharmacovigilance efforts primarily monitor the frequency of ADRs, and typically does not focus on the potential and differential impact of those ADRs on patients.

Analytical methods have shifted from qualitative to quantitative as the availability of data, metrics, and computing power continues to increase. The existing categorical definitions of ADR severity limit the ability of researchers and regulators to apply quantitative methods to regulatory and pharmacovigilance efforts. For instance, tracking the regulatory performance of drug safety efforts is primarily done through categorical analysis of ADR case outcomes and drug label changes. The creation of a quantitative numerical scale providing a relative severity score for each ADR would enable these analyses to leverage the powerful tools of quantitative decision and utility theory.

Pioneering work by Tallarida et al. examined the ability to create a continuous ADR severity scale[7]. 53 physicians were interviewed and asked to estimate acceptable probabilities in a set of scenarios specifying risk benefit tradeoffs. These probabilities were then used to define a set of equations and the expected benefit to the patient in each scenario. Solving the system of equations resulted in numerical severity weights that were relative to the lowest severity ADR. The effort took ~45 minutes per interview and resulted in relative scores for 7 ordinal categories of ADR severity.

To expand this effort, Gottlieb et al. expanded this ADR severity ranking task using the crowdsourcing platform Amazon Mechanical Turk[8]. Individual workers were presented with a pair of ADRs, as well as a link to more information on those ADRs, and were asked to select the ADR they perceived to be more severe. Over 146 person days, 2,589 workers produced severity comparisons for 126,512 ADR pairs composed from 2,929 unique ADRs. A linear programming algorithm was used to create a unified ADR severity ranking. To validate the ranking, ADR severities were correlated against ADR case outcomes from the Food and Drug Administration’s Adverse Event Reporting System (FAERS). The highest Spearman correlation, *ρ =* 0.53, was with proportion of cases resulting in death. This effort drastically increased the number of ADRs included in the severity ranking, but was costly with regards to human time, labor, and money.

Recent work in natural language processing has resulted in numerous methods for the creation of vectorized word representations (also known as “embeddings”) that capture semantic meaning in a dense numerical representation[9,10]. These methods rely on the distributional hypothesis, that word meaning is captured by the contexts in which a word appears[11]. The practical implication of the distributional hypothesis is that training a model to predict word context (i.e. co-occurring word pairs) results in model weights that capture the meaning of the word. These model weights can then be utilized as a numerical representation of a given word.

Word embeddings learned on social media datasets have been deployed for pharmacovigilance before, but not for the purpose of exploring ADR severity[12]. Using a social media corpus generated by the general public, especially on pseudo-anonymous social media platforms like Reddit, can capture meanings that reflect people’s unfiltered experiences of and opinions about health and disease[13,14]. Previous research on Twitter data annotated samples of tweets for personal medication intake versus individual’s simply mentioning a drug [15,16]. Together, the two studies found ~40% of tweets mentioning a drug indicated the individual tweeting was possibly personally taking that medication. Here we are focused on ADRs, but it is likely that individual’s discussing ADRs on social media have more direct experience with the ADRs, either by being directly affected by the ADR or being informed about the ADR experience by a close relation than individuals selected from a pool of crowdworkers. Word embeddings trained on a corpus generated by the general public can then be leveraged as a metric for public opinion in a similar fashion as previous crowdsourced approaches. We use the RedMed embeddings trained on over 580 million health-enriched Reddit comments from >10 million users[17]. These numbers dwarf the number of votes gathered in a typical crowdsourcing experiment and are therefore potentially more indicative of a representative populations’ perception of ADRs than traditional survey-based methods.

In this paper, we use publicly available word embeddings and a network-based label propagation method to estimate the severity of 12,198 ADR concepts from the MedDRA terminology. The resulting Severity of Adverse Events Derived from Reddit (Saedr) scores are validated against human rankings, as well as FAERS case outcomes. We use System Organ Classes (SOCs) and other groupings within MedDRA to examine how Saedr severity scores differ at various levels of abstraction and within ADR categories. We use Saedr scores to compare severity between ADRs in different sections of drug labels, ADRs with disproportionate rates between sexes, and ADRs discovered at different stages of drug development. We combine Saedr scores with frequency information from SIDER-derived drug labels to generate drug-specific aggregate side effect severity scores[18]. Saedr scores enable new analyses that were not previously possible with existing categorical classifications.

## Methods

### 1. Data sources and preparation

#### 1.1 ADR terms and phrases

We sourced our ADR phrases and their synonyms from “lower level terms” (LLTs) within the Medical Dictionary for Regulatory Activities Terminology v22 (MedDRA)[19]. We filtered terms based on their semantic types within the UMLS Metathesaurus[20] and retained terms of the following semantic types: “Disease or Syndrome” (T047), “Finding” (T033), “Neoplastic Process” (T191), “Injury or Poisoning” (T037), “Pathologic function” (T046), “Sign or Symptom” (T184), “Mental or Behavioral Dysfunction” (T048), and “Congenital Abnormality” (T019).

#### 1.2 Word embedding model

We utilized the word embedding model from the RedMed project[17]. This model was selected as it was directly optimized for medical term similarity, was trained on a corpus of Reddit comments generated by the public and preprocessed to maximize the inclusion of ADR terms. ADRs that were a phrase, such as “abdominal pain”, were represented using the average embedding of all terms within the phrase. We found the use of average embeddings, e.g. “abdominal” + “pain”/2, produced better results than the use of embedded phrases, e.g. “abdominal_pain”.

#### 1.3 Gottlieb severity data

We used the crowdsourced severity estimates included within the supplement of the work by Gottlieb et al., there were a total of 2,929 ADRs with rank scores[8].

#### 1.4 FDA Adverse Event Reporting System data

All drug adverse event data was downloaded in JSON format from the openFDA website (https://download.open.fda.gov/drug/event/), which includes data from both the Legacy Adverse Event Reporting System (LAERS), as well as FDA Adverse Event Reporting System (FAERS). Data was retrieved on August 08, 2020 and included adverse event case reports up until June 30, 2020. Drug reactions were normalized to MedDRA v22. We filtered to unique caseids using the provided duplicated flag to remove duplicate reports and case reports that did not originate in the United States. Case outcomes were normalized according to the following schema: “Death”: {“Death”}, “Serious Outcome”:{“Life-Threatening”, “Hospitalization”, “Other Serious”, “Required Intervention”}, “Disability”: {“Congenital Anomaly”, “Disability”}. We created an additional outcome category denoted “No Outcome” for cases with no reported outcome. We felt the need to create this category as FAERS only allows for the reporting of serious case outcomes and there is information in the absence of a serious outcome being reported. Outcome proportions for each ADR were calculated by dividing the number of cases reporting that ADR for a particular outcome by the total number of cases reporting that ADR.

#### 1.5 FAERS severity rankings

For the ranking of ADRs based on FAERS data we ranked ADRs based on their marginal likelihood to be included in a case with Death or Serious Outcome as the reported case outcome. This was calculated by dividing the number of cases with the ADR that resulted in Death or Serious Outcome by the number of cases without the ADR that resulted in those outcomes.

### 2. Semi-supervised severity propagation

Given a word embedding model and a set of initial, potentially noisy, seed word labels for the severe and benign categories, we seek to propagate severity information over the rest of the vocabulary similar to the sentiment propagation methods of Hamilton et al.[21]. A graphical overview of the method is provided in Figure 1.

**Figure 1:**
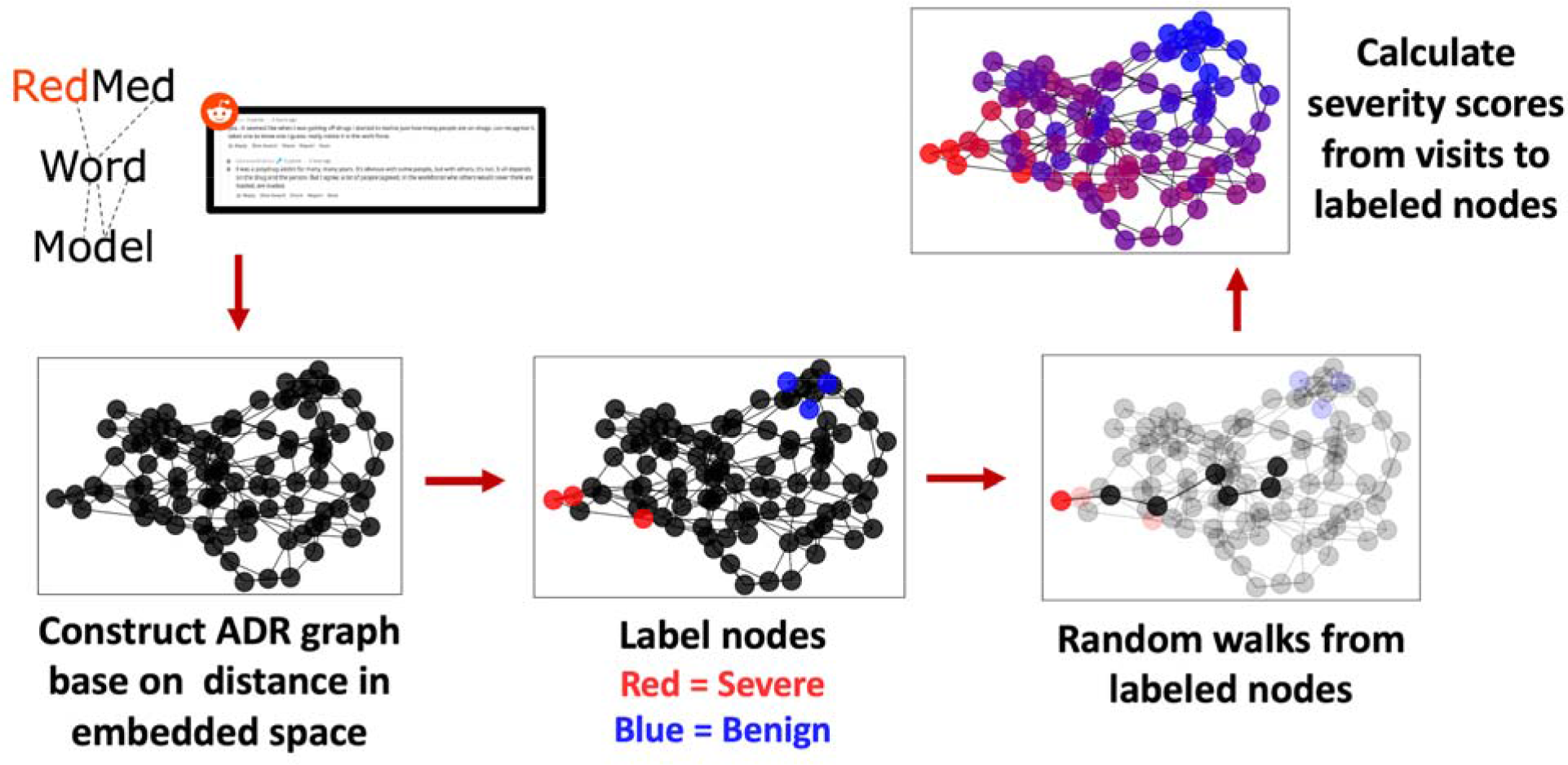
Overview of network method for estimating ADR severity from word embeddings. Word embeddings for 28,113 ADR terms and phrases were extracted from the RedMed word embedding model. A network is constructed based on nearest-neighbors in the embedding space. A subset of nodes are labeled as Severe or Benign ADRs and random walks from those labeled nodes are conducted. Saedr scores for each ADR are calculated based on the relative number of encounters in random walks initiated from severe versus benign nodes.

#### 2.1 Lexical network creation

The network is constructed by connecting words (nodes) with edges to their ***k*** nearest-neighbors based on the cosine distance between their embeddings. The corresponding edge weight is set to the cosine similarity of the two words *(i.e.* edge weight increases with term similarity).

#### 2.2 Seed selection from terminology

An initial set of ranked seed terms are labeled. Biomedical terminologies and ontologies often contain many terms that are highly similar to each other. To reduce the effect of lexical similarities on label propagation, we filter the benign and severe seed terms based on the real minimal edit distance between terms in each list. Only terms that have a distance > 0.5 from a higher ranking term are included in the list, resulting in a reduced list of lexically distinct seed terms. Both benign and severe seed term lists are truncated to the minimum seed term list size of both seed sets.

For instance, if a severe seed term list had the following items in rank order, [“death”, “cardiac arrest acute”, “cardiac arrest”] and the benign seed term list [“dry skin”, “yawning”, “cold sweat”]. After lexical filtering, the severe seed term list would be [“death”, “cardiac arrest acute”] and the benign seed term list would remain unchanged. The final seed term lists would be truncated to the minimum seed term list size, resulting in severe seed term list [“death”, “cardiac arrest acute”] and a benign seed term list [“dry skin”, “yawning”].

#### 2.3 Random walks from seed nodes

Given a network and a set of seed nodes, we modified code from node2vec[22] to perform 5,000 weighted random walks, of length 200, from each seed node. We selected the number of random walks to ensure that all nodes within the graph were visited a non-zero number of times. We note that this value needs to be empirically determined and will likely change based on the size and structure of the network.

#### 2.4 SAEDR Score calculation

The Saedr score of a given node *u* is calculated using the following formula:

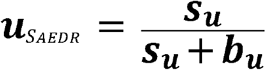

Where *s_u_* and *b_u_* are the respective number of times that node *u* is encountered in a random walk from a severe node or a benign node. If node*u* is contained within one of our seed nodes sets then self-visits during the random walks are excluded from the calculation. To increase the robustness of these score estimates we perform 10,000 iterations of bootstrap sampling[23] of the random walks and the average of these bootstrap estimates is calculated. Saedr scores from multiple seed sets are averaged at the PT level and the final combined scores are normalized to the interval [0,1].

### 3. Hyper-parameter tuning for severity propagation

We randomly split the crowdsourced severity data into training (75%, n=1,778) and test (25%, n=591) sets with 284 ADRs dropped due to mapping. We performed a grid search over the number of neighbors used to construct the lexical network {2, 5, 10, 15, 20, 25, 30} and the percent of nodes to label for the severe and benign seed nodes {2, 5, 10,15, 20, 25}. For example, an individual run would use the 25 nearest-neighbors and 5% at each end of the training severity rank (i.e. the top 5% most severe and bottom 5% least severe ADRs). We ultimately selected the parameters with the highest Spearman correlation with the training data.

### 4. ADR discovery group analysis

We analyzed the Saedr score distributions of several different categories of ADRs.

#### 4.1 ADRs included in a Boxed Warning

We downloaded counts for the number of appearances of ADRs in the Boxed Warning section of drug labels from the supplement of Wu et al.[24]. ADRs that appear on a drug label, based on their appearance in SIDER, that were not included in the list provided by Wu et al. were considered to not have appeared in a Boxed Warning section [18].

#### 4.2 ADRs with disproportionate reporting between sexes

A set of ADRs shown to have sex differences were identified from the supplement of Chandak et al.[25]. We filtered out ADRs considered Borderline, (mean log(ROR) < 0.4 and mean log(ROR) > −0.4), as defined in the original work by Chandak. Their effort reported ADRs at the Higher Level Group Term (HLGT) level within MedDRA, while our severity estimates are generated at the Preferred Term (PT) level. There are many PT ADRs per HLGT, so all PTs within the specified HLGT terms were averaged to create a single Saedr score for that HLGT. We did not use the drug information except to note distinct instances of sex specific HLGTs (i.e. HLGTs associated multiple times with different drugs each time).

#### 4.3 ADRs discovered at different stages of drug development

ADRs discovered at the clinical trial stage were identified based on their inclusion in a drug label on SIDER with a non-postmarket ADR frequency[18]. Postmarket ADRs were identified as those included on drug labels with a postmarket frequency and no clinical trial frequency information. OffSides and TwoSides ADRs were discovered from FAERS data looking at disproportionate reporting after correcting for the effects of demographics and other case information[26]. We consider both OffSides and TwoSides ADRs as discovery stage postmarket ADRs, as they have not yet been included in drug labeling.

### 5. Drug RIsk Profile scores

Drug RIsk Profile (Drip) Scores were calculated based on our Saedr score and the frequency of each ADR for a given drug. The Drip score for a given drug is the sum of the severity multiplied by the frequency for each ADR on the drug label. Drug ADR frequency information was retrieved from the SIDER resource[18]. In situations where multiple ADR frequencies were indicated, 1,000 frequency estimates were sampled from a uniform distribution with a lower bound at the minimum reported frequency and an upper bound at the maximum reported frequency. When there was no frequency information we sampled 1,000 estimates from a uniform distribution over the interval [0.001, 0.01]. The final Drip score is the average score across 1,000 samples.

## Results

### Network statistics

We used the ADR terms and phrases from MedDRA v22.0 LLTs[19] as the initial lexicon for the lexical ADR network. While FAERS uses MedDRA to encode ADRs, versioning differences over the years resulted in some FAERS ADRs not being included in MedDRA. We were able to generate embeddings for 92% (2,450/2,653) of unique crowdsourced ADRs, 100% (14,045/14,045) of unique FAERS ADRs, and 69% (28,113/40,905) of all filtered MedDRA terms. This resulted in a final lexical network with 28,113 nodes, representing 12,198 MedDRA PTs.

### SAEDR score performance

We compared our severity estimates using crowdworker ranked seeds, FAERS ranked seeds, and the average severity estimate across the two seed sets to two different ADR rankings. (1) A held-out test set of crowdworker ranked ADRs (n=591) and (2) ADRs ranked by case outcome statistics in the FAERS database.

The crowdworker seeded severity estimates had the highest training performance with a 25 nearest-neighbors graph and using 10% of the most and least severe ADRs as seeds. The Spearman correlation with the crowdworker test set was 0.747. Spearman correlations for events with at least 100 reports in FAERS were 0.595, 0.616, and −0.732 for death, serious outcome, and no outcome, respectively (Supp. Fig 1, Supp. Fig 2).

The FAERS seeded severity estimates had the highest training performance with a 10 nearest-neighbors graph and using 10% of the most and least severe ADRs as seeds. The Spearman correlation with the crowdworker test set was 0.587. As no information from the crowdworker rankings was used to select these seeds, we can also report the Spearman correlation with the entire set of crowdworker ADRs, 0.765. Spearman correlations for events with at least 100 reports in FAERS were 0.509, 0.557, and −0.656 for death, serious outcome, and no outcome, respectively (Supp. Fig 1, Supp. Fig 2).

The Saedr score, the average of the two severity estimates, had a Spearman correlation with the crowdworker test set of 0.735 (Figure 2B). Spearman correlations for events with at least 100 reports in FAERS were 0.595, 0.633, and −0.748 for death, serious outcome, and no outcome, respectively (Supp. Fig 1, Supp. Fig 2).

**Figure 2:**
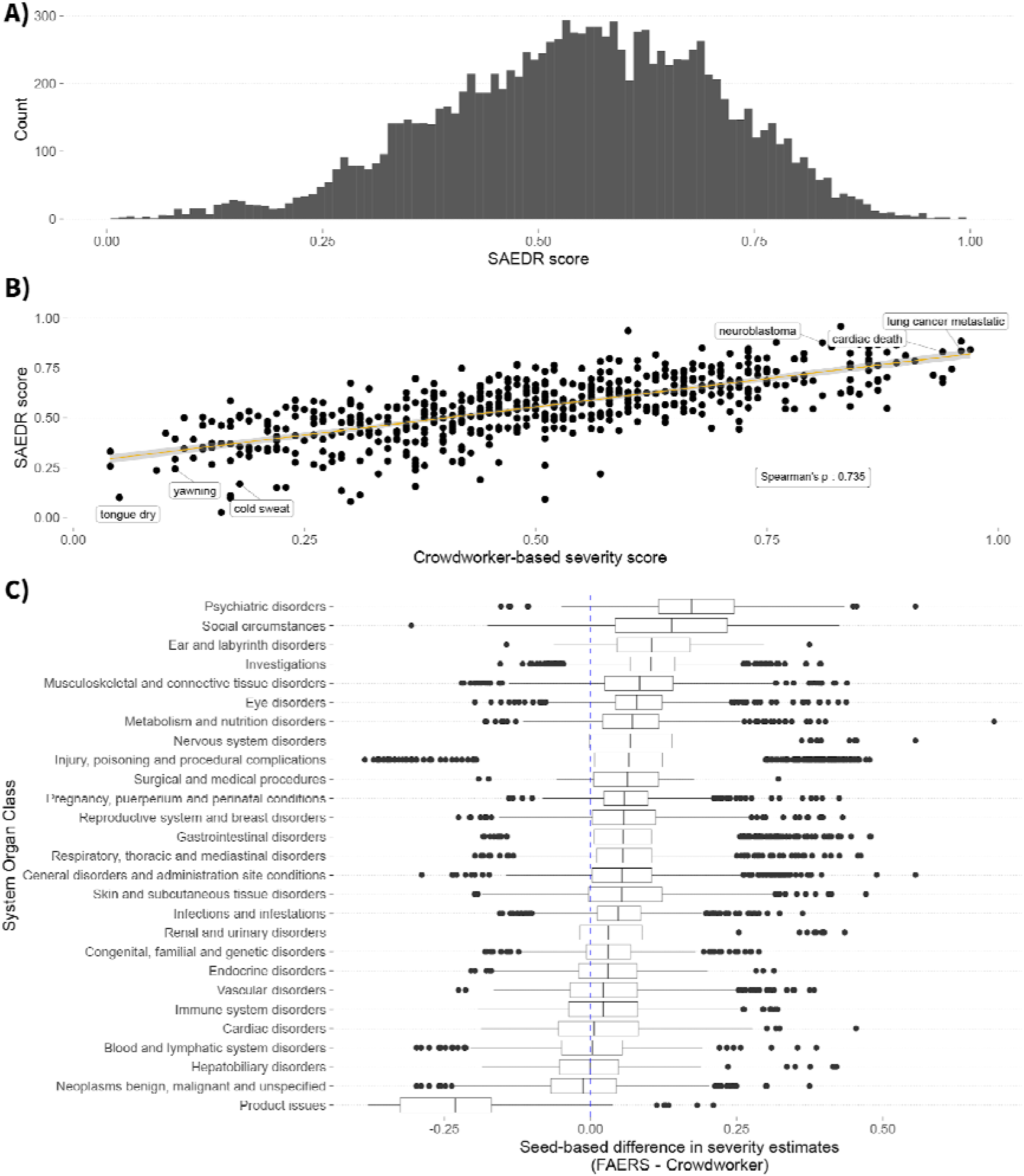
Comparison of SAEDR scores to crowdworker severity estimates. **A)** Histogram of Saedr scores for 12,198 ADRs. **B)** Crowdworker severity estimates (x-axis) versus Saedr scores (y-axis) for a test set of 591 ADRs. Saedr scores show a strong Spearman correlation, *ρ* = 0.735, with the hum**an** crowdworker rankings. We note that this correlation is greater than the inter-rater correlation, *ρ* = 0.71, reported in the original crowdsource effort by Gottlieb et al. A select set of the least and most severe ADRs based on the SAEDR score have been annotated. **C)** Differences between severity estimates seeded with FAERS rankings versus those seeded with crowdworker rankings (x-axis) for different SOC groups (y-axis). Dashed blue line indicates where severity would be the same for both estimates. We observe the biggest difference in severity for “Psychiatric disorders”, “Social circumstances”, and “Product issues”. With the two former categories having increased severity from the FAERS seeds and the latter category having markedly lower severity from the FAERS seeds.

### Seed ranking comparisons

We examined the effects of seeding on the severity estimates for individual ADRs. Most ADRs were not substantially changed by the use of different seeds. We aggregated ADR severity differences at the MedDRA System Organ Class level and found that the FAERS seeds led to increased severity estimates for “Psychiatric Disorders” and “Social Circumstances” (Fig. 2B). The most drastic shift in values was the decrease in severity of “Product Issue” related ADRs using FAERS seeds relative to their severity when using crowdworker based seeding.

### System Organ Class severity rankings

We compared the severity of ADRs based on MedDRA System Organ Class groupings as a qualitative evaluation (Figure 3A). We find ADRs related to cancer (Neoplasms), cardiac, and liver (Hepatobiliary) to be the most severe, high Saedr scores, while skin, general disorders, and product issues are considered the least severe, low Saedr scores. We performed a qualitative examination of ADRs at various other levels. We compared severity distributions between ADRs describing benign versus malignant neoplasms (Supp. Fig. 3). A one-sided t-test found malignant neoplasms to be significantly more severe than benign neoplasms based on their Saedr scores (p < 0.001). We found there to be wide ranges of severity within the different levels for MedDRA ADR term groupings (Figure 3B, C, and D). We note that among ADRs in the “Cardiac disorders” HLGT, cardiac neoplasms have the highest Saedr scores, while signs and symptoms of cardiac disorder have the lowest relative Saedr scores (Figure 3B). When examining the level of individual ADRs within the “Heart Failure Signs and Symptoms” we find ADRs we would judge more severe ranked higher than those that are more general and considered to be less severe.

**Figure 3.**
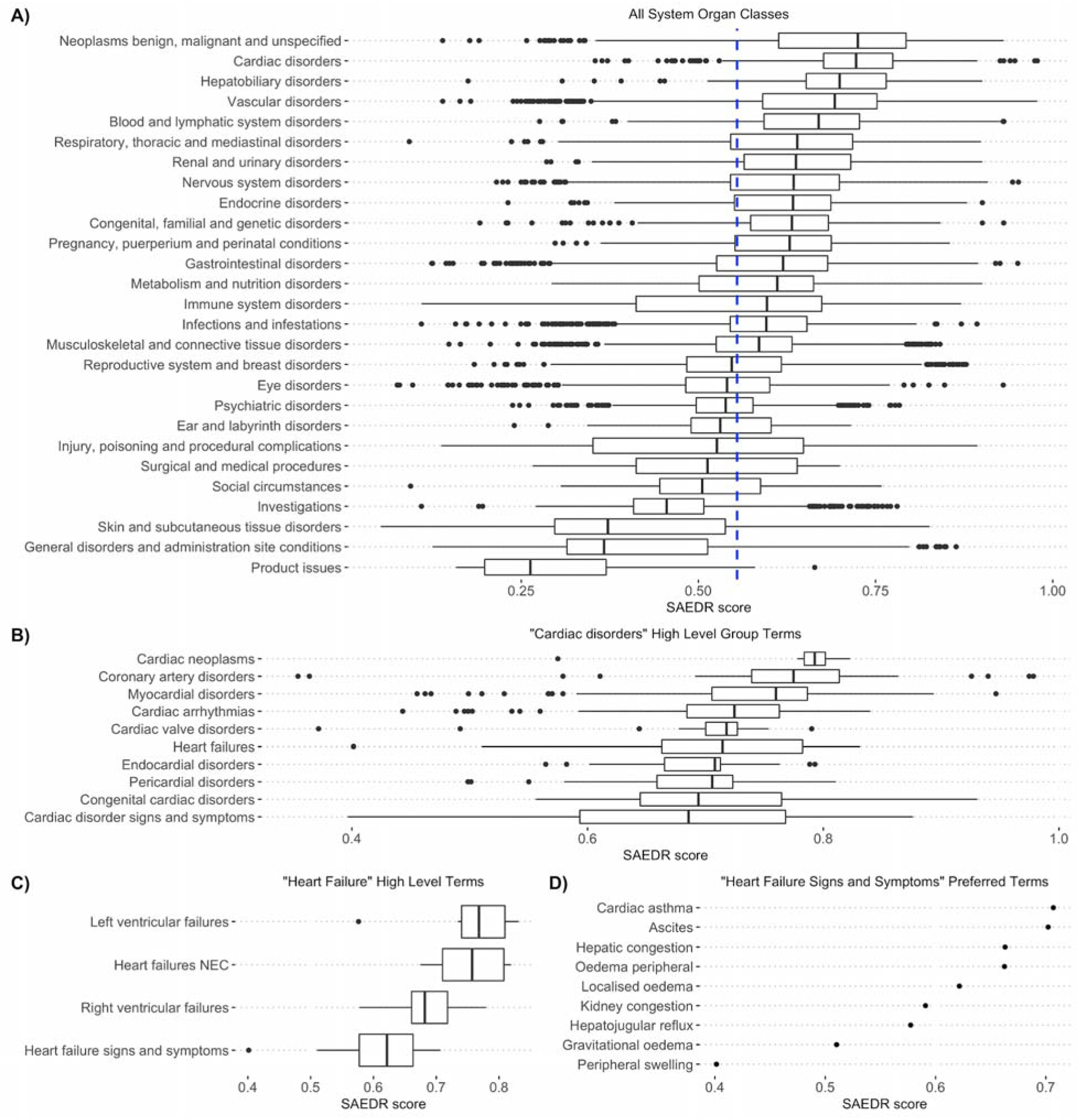
Severity of ADRs at different group resolutions. A) Saedr score (x-axis) for 11,981 ADRs with SOC groups (y-axis). Dashed blue line indicates overall median Saedr score of 0.554. Saedr score distributions indicate that cancers and cardiac related side effects are considered the most severe, with product issues being the least severe. B) Severity distributions of MedDRA High Level Group Terms (HLGT) within the “Cardiac Disorders” SOC. There are large differences in severity within these term groups. C) High Level Terms (HLT) within the “Heart Failure” HLGT show a tighter distribution of severity, with heart failure signs and symptoms being the least severe within that group. D) Looking at individual side effects within the signs and symptoms of heart failure, we see cardiac asthma and ascites, accumulation of fluid in the peritoneal cavity, as the most severe symptoms, while peripheral swelling is considered the least severe.

### Severity of ADRs grouped by labeling section, sex effects, and time point of discovery

Based on our estimate, we calculated that ADRs that have never been included in a Boxed Warning had a median Saedr score of 0.538, while ADRs that were included in at least 1 Boxed Warning had a median Saedr score of 0.624. We found that among ADRs that have appeared on a drug label, those that have been included in at least 1 Boxed Warning were significantly more severe (p < 0.001) than ADRs that have never been included within a Boxed Warning section, based on a one-sided t-test (Figure 4A).

**Figure 4.**
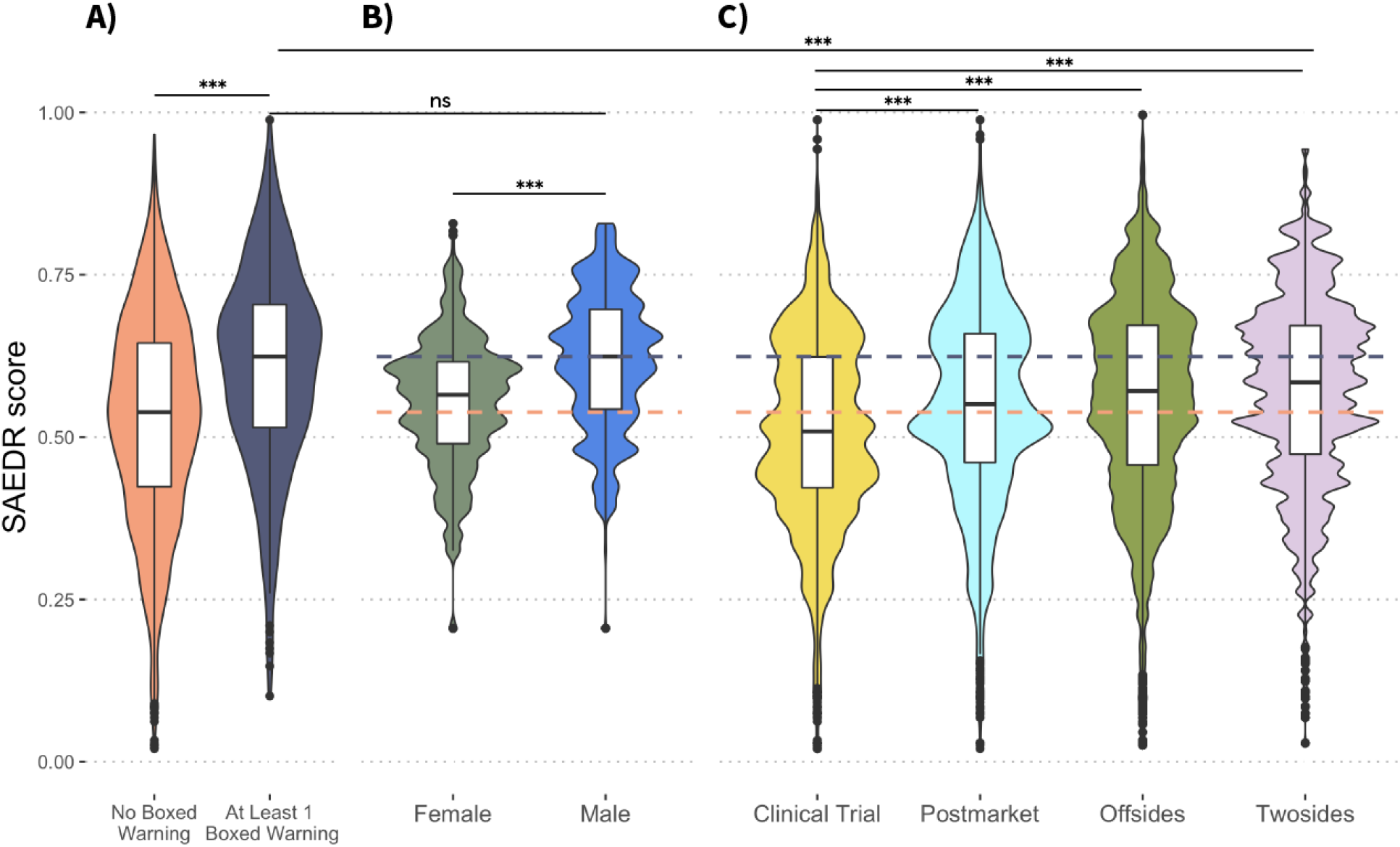
Differences in ADR severity between ADR groupings and discovery periods. ADR groups (x-axis) vs Saedr scores (y-axis) Grey dashed line indicates median severity, 0.624, of ADRs that have been included in a boxed warning. Orange boxed line indicates median severity, 0.538, of ADRs that appear on a drug label but have not been included in a Boxed Warning. **A)** ADRs that were listed as a black box warning at least once (n = 356) were significantly more severe than ADRs that have not appeared in a black box section (n = 3,305). **B)** ADRs that disproportionately affect males (n = 56,405) are significantly more severe than those that disproportionately affect females (n = 50,801). There was no significant difference in severity between ADRs included in black box warnings and ADRs that disproportionately affect males. **C)** ADRs discovered postmarket (n = 11,506) are significantly more severe than those discovered in clinical trials (n = 35,450). ADRs identified postmarket via OffSides (n = 350,631) and postmarket polypharmacy ADRs identified via TwoSides (n = 4,210,513) are significantly more severe than those discovered in clinical trials. The severity of all postmarket ADR groups is significantly less severe than ADRs that have appeared in a black box warning.

We found ADR HLGTs that are disproportionately reported for males were significantly more severe (p < 0.001) than those disproportionately reported for females (Figure 4B). HLGTs disproportionately experienced by males were not significantly different (p > 0.05) from the severity of ADRs that have been included in at least 1 Boxed Warning, based on a one-sided t-test.

We examined the severity of ADRs that were found during the clinical trial versus those discovered postmarket, based on SIDER label annotations. We found ADRs discovered postmarket to have significantly higher severity (p < 0.001) than ADRs discovered in clinical trials, based on a one-sided t-test (Figure 4C). We compared the severity of the clinical trial ADRs to those discovered postmarket using FAERS data. Tatonetti et al. identified two differents sets of ADRs, OffSsides is a set of ADRs that were disproportionately reported for a drug, while TwoSides is a set of ADRs that were disproportionately reported for a pair of drugs being taken concurrently, after correcting for case demographic and other information. The severity distribution of ADRs in both OffSsides and TwoSides were significantly higher (p> 0.001) than those discovered in clinical trials, based on a one-sided t-test (Figure 4C). We found ADRs associated with polypharmacy, via TwoSides, to have the highest severity of the postmarket ADRs. All postmarket ADRs were significantly lower in severity than the Boxed Warning ADRs, based on a one-sided t-test.

### DRIP score analysis

We calculated Drip scores for 968 drugs with SIDER label data, with a resulting median Drip score of 0.439. The Spearman correlation between drugs ranked by the proportion of FAERS cases with that drug as the primary suspect that resulted in death and drugs ranked by our Drip scores was 0.377.

We analyzed Drip score distributions by Anatomical Therapeutic Chemical (ATC) for the subset of drugs with an ATC designation (Figure 5A). Antineoplastics and antiepileptics were the drug classes with the highest Drip scores, indicating drugs in these classes have severe ADR profiles. Drugs used in the management and treatment of diabetes and urological issues had the lowest Drip scores, indicating these drugs are relatively safe and have less severe side effect profiles. We examined individual drugs within the opioid class and found fentanyl to have a markedly higher Drip score than other drugs in the same class (Figure 5B). Drugs within the statin class were primarily below the overall median Drip score (0.439), with only atorvastatin being markedly above the median Drip score (Figure 5C).

**Figure 5.**
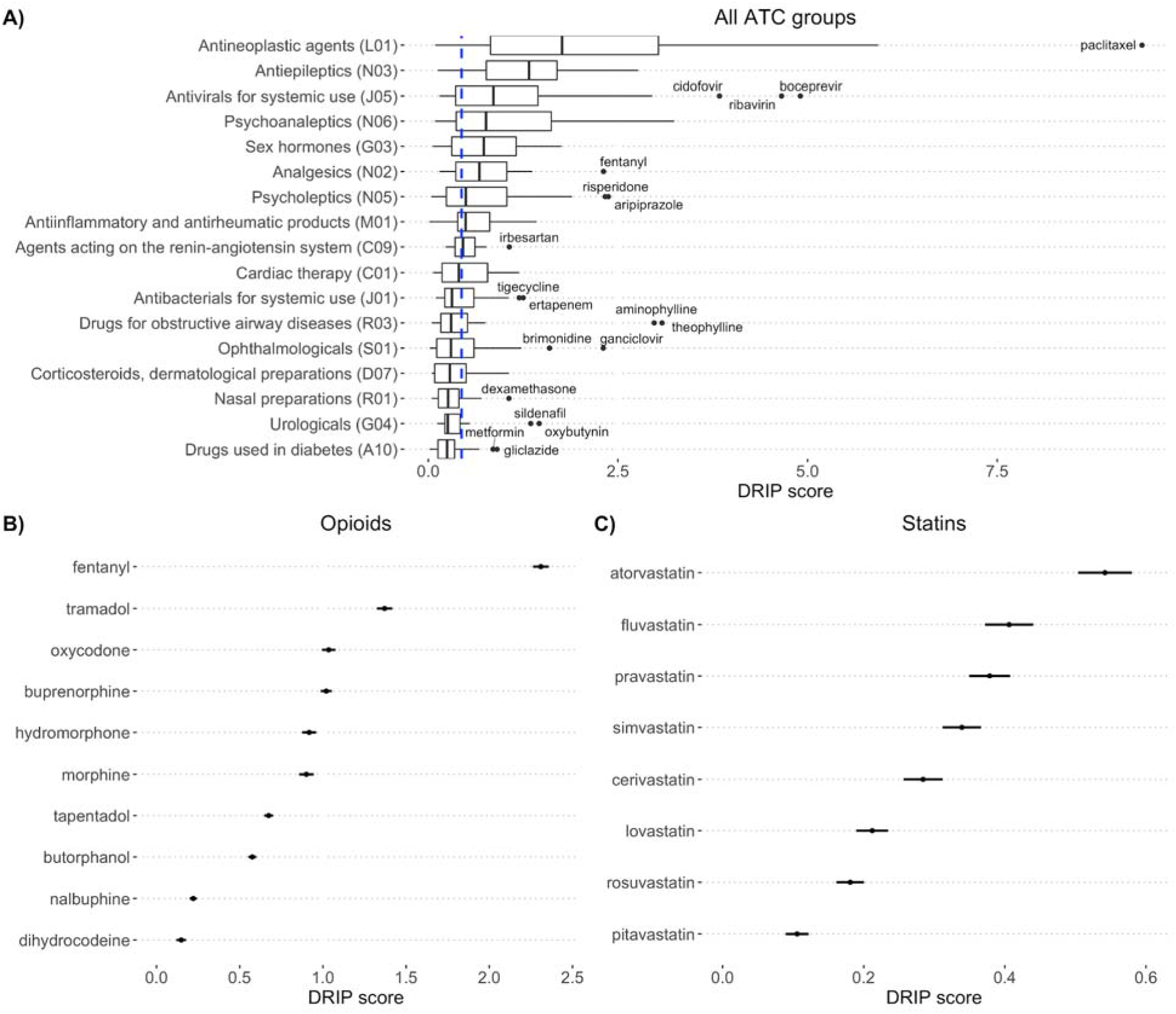
Drug RIsk Profile (**Drip**) scores calculated using side effect severity and frequency. A) Drip scores (x-axis) for ATC groups (y-axis) ordered by median severity with outliers labeled. Dashed blue line indicates the median Drip score of 0.439 across all drugs. **B)** Drip scores (x-axis) for opioid class drugs (y-axis), as denoted by the ‘N02A’ ATC group. Fentanyl, an extremely potent synthetic opioid is ranked first as having the most severe side effect severity. **C)** Drip scores (x-axis) for statins (y-axis), as denoted by the ‘C10AA’ ATC group. Atorvastatin, the strongest statin in broad use, is ranked to have the most severe side effect severity, while lower efficacy statins are ranked as having less severe side effect severities.

## Discussion

In this work, we present Saedr scores, quantitative estimates of the severity of 28,113 MedDRA ADR terms. Saedr scores have strong correlations with both crowdworker based severity estimates and real-world ADR case outcome statistics. We generated these estimates using a network-based label propagation approach that required only a small percentage of the terms to be labeled. This effort demonstrates the feasibility of using distantly supervised techniques, such as label propagation, to generate quantitative values for medical concepts.

Our approach enabled us to increase the number of ADRs with quantitative severity estimates from 2,929 to 28,113, an almost 10-fold increase, while minimizing the human time involved in generating the estimates. Our severity estimates can be updated routinely based on recomputation of the embeddings and re-seeding of the reference severe/benign ADRs. We note that our severity estimates had a higher correlation with the aggregate crowdworker rankings, *ρ =* 0.735, than the crowdworker rankings had between the three replicates in the original study, *ρ =* 0.71. Additionally, our estimates had a greater correlation with real-world outcomes, such as FAERS cases resulting in death, *ρ =* 0.595, than the crowdworker rankings, *ρ =* 0.53. Thus, our severity estimates track with both human judgements of ADR severity, as well as real-world outcomes of consequence to patient health.

We found our Saedr scores had a strong negative correlation, *ρ =* −0.748, with no case outcome being reported in FAERS. We assigned a “No outcome” label to FAERS cases where none of the available, but relatively severe, case outcomes had been reported. We take this as an indication that FAERS cases lacking outcomes may be a result of cases having an outcome below the level of seriousness that is reportable in FAERS. FAERS case outcome completeness could potential be improved with the creation of a lower severity category for case outcome documentation.

In an effort to compare the impact of using different methods for selecting the initial severe and benign ADR seed terms for label propagation, we compared crowdworker versus real-world outcomes-based seed selection. We found that both methods resulted in consistently high severity scores for ADRs in the highest severity SOC groups, such as neoplasms and cardiac disorders. We observed the largest differences in, and product issues SOC groups. Real-world outcome-based seeds increased the severity of psychiatric disorders, social circumstances and decreased the severity of product issues. The importance and impact of psychiatric disorders may still not be apparent to those not in the medical profession. Thus, it may be that crowdworkers did not perceive mental illness to be as severe as physical illnesses, leading to a bias in the rankings. While using real-world outcome-based rankings to select seeds, led to more psychiatric disorders being included in the seed terms for the severity propagation. Similarly, the shifts in social circumstances and product issues is likely due to their limited presence or absence within the original crowdworker rankings. As only 7 ADRs in the social circumstances category were included in the 2,929 ADRs ranked by crowdworkers, and no “product issues” ADRs were included in that ranking. This absence resulted in no ADRs from these groups being included in the initial seed set of labeled ADRs for the crowdworker seeded method run, likely limiting the ability of the label propagation to accurately estimate the severity of these ADRs.

We found the overall severity ordering of the SOC classes based on our Saedr scores to make intuitive sense. We were surprised by the ability of the label propagation approach to adjust severity estimates based on particular key terms. For instance, the ADR terms for cancer (“Neoplasm”) ADRs are relatively similar on a lexical basis, but individual modifiers such as malignant and benign resulted in significantly different Saedr scores for the respective groups of ADRs. Similarly, examining groups of ADRs at different MedDRA hierarchy levels demonstrated relative severity estimates that were sensical to us, such as “heart failure signs and symptoms” having lower severity than the different types of actual heart failure.

We found a significantly higher estimate of severity among ADRs that had been included in a Boxed Warning section versus those that had not. This offers further validation of the Saedr scores, as they are in agreement with past regulatory and drug labeling decisions. Our comparison of HLGTs with disproportionate rates between sexes, found male-associated ADRs to be more severe. Notably, the Saedr scores of male associated ADRs were not significantly different from the Saedr scores of ADRs that have been included in a Boxed Warning. This is in agreement with previous work that has highlighted the relatively higher severity of ADRs experienced by males[27,28].

We found that ADRs discovered postmarket are generally more severe than those discovered in the clinical trial based on our Saedr scores. Indicating that postmarket surveillance and ongoing regulatory discussions of risk-benefit tradeoffs for particular drugs are a necessity to keep the public safe [29]. These findings are not new, as a recent study examining internal FDA data on drug labeling changes found that 35% of ADRs added to drug labels were added to the Boxed Warnings and Warnings and Precautions sections[30]. Postmarket ADRs involving more than one drug, as identified by TwoSides, were among the highest severity of the postmarket ADRs indicating the increased risks associated with polypharmacy. This finding highlights the need for further research into the safety of polypharmacy, as approximately half of individuals prescribed a prescription drug are prescribed more than one concurrent medication[31].

One of the original aims of the pioneering work by Tallarida et al. was to create quantitative risk scores for individual drugs. Our Drip scores are an attempt to do so, we combined ADR severity with frequency information to generate a numerical estimate of a drug’s risk profile. We were able to generate Drip scores for 968 drugs with ADR frequency data from SIDER. Our Drip scores have a modest Spearman correlation, *ρ =* 0.377, with FAERS cases that resulted in death, where that drug was the primary suspect. We interpret this as an indication that our Drip scores track with real-world outcomes and capture signal related to a drugs safety profile. A qualitative evaluation of Drip score distributions for ATC groups makes intuitive sense, with antineoplastic drugs having the most severe side effect profiles, while diseases that are known to have safe and effective treatment options, such as diabetes, are ranked among the safest.

We present ADR severity estimates that track with both real-world outcomes and human perception, but these estimates are still limited. We used word embeddings trained on Reddit data, but other social media data could improve the severity prediction model. We used RedMed, as the model was publicly available and contained many ADR terms of interest to this effort. However, Reddit as a social media platform is skewed towards males and young people, who may discuss ADRs in different ways than the general population. Learning new embeddings from social media corpora generated by patient groups most at risk for a given set of ADR experiences or from biomedical literature might improve model performance and address issues with demographic model biases.

Our results indicate that males reported more severe ADRs based on sex-specific ADRs identified by Chandak et al. Other research efforts have reported similar findings based on ADR reporting databases[27], but there are potential sources of bias that could have impacted this result. The data underlying the sex-specific ADRs identified in Chandak et al. was derived from FAERS. Their method focused on the analysis of a sex-balanced cohort of case reports, with cohort creation via propensity-score matching of individuals. Propensity score matching can correct for factors included in the model, but it is unable to correct for underlying biases in reporting. It has been documented that there are higher rates of ADR reporting for female patients than for males, while males tend to have more severe outcomes reported[28]. It’s likely that males are enriched for severe ADRs due to this reporting bias and ADR reports should not be conflated with all ADRs experienced by individuals due to the low rate of ADR reporting[32]. Another limitation is that sex-specific ADRs were only reported at the HLGT level, resulting in Saedr scores that were an average of all PTs within the HLGT. Figure 3 highlights the large range of severities even within a particular HLGT. It is possible that an analysis of sex-specific ADRs at the PT level would result in a different finding. Overall, while ADRs disproportionately reported in men are more severe than those disproportionately reported in women, more research is needed to reach a complete understanding of sex-specific differences in ADR severity.

While we capture some nuance in ADR severity, such as benign versus malignant neoplasms, we only created Saedr scores for the ADR terms contained in MedDRA. However, while exploring the RedMed word embeddings, we saw individual ADR terms were often contained in phrases indicating modified severity. For instance, a report of stomach pains might be modified with an adjective such as excruciating or mild. While MedDRA does not contain an intensity scale component for its terms, capturing this information from patients might further enable risk-benefit tradeoff calculations. Approaches for general word sentiment assignment, similar to the one used by Hamilton et al., could be repurposed here to assign severity to ADR modifier terms[21].

Our Drip scores would be improved by more comprehensive estimates of ADR frequencies for drugs. The majority of ADR frequencies extracted by SIDER are from the clinical trial and are affected by dosage and underlying health of the clinical trial population, both of which are only sometimes reported. Frequency estimates for postmarket ADRs are essentially non-existent and many challenges exist to accurately estimating their frequency. We view comprehensive ADR frequency estimates as necessary for the accurate assessment and quantification of drug risk profiles.

In summary, we demonstrate that lexical networks and label propagation can be used quantitatively to estimate the severity of medical conditions. We show distributions of ADR severity among different groups of conditions, different groups of patients, and different ADR discovery time points. We combined our Saedr scores with available ADR frequency data to generate quantitative scores of drug risk profiles and examined the distribution of the resulting Drip scores. Our results (and future improved estimates) enable new quantitative analyses within the field of pharmacovigilance. To this end, we provide the complete set of Saedr scores and Drip scores in the supplement of this paper.

## Supporting information

SAEDR Scores

DRIP Scores

---

**Supplementary Figure 1.**
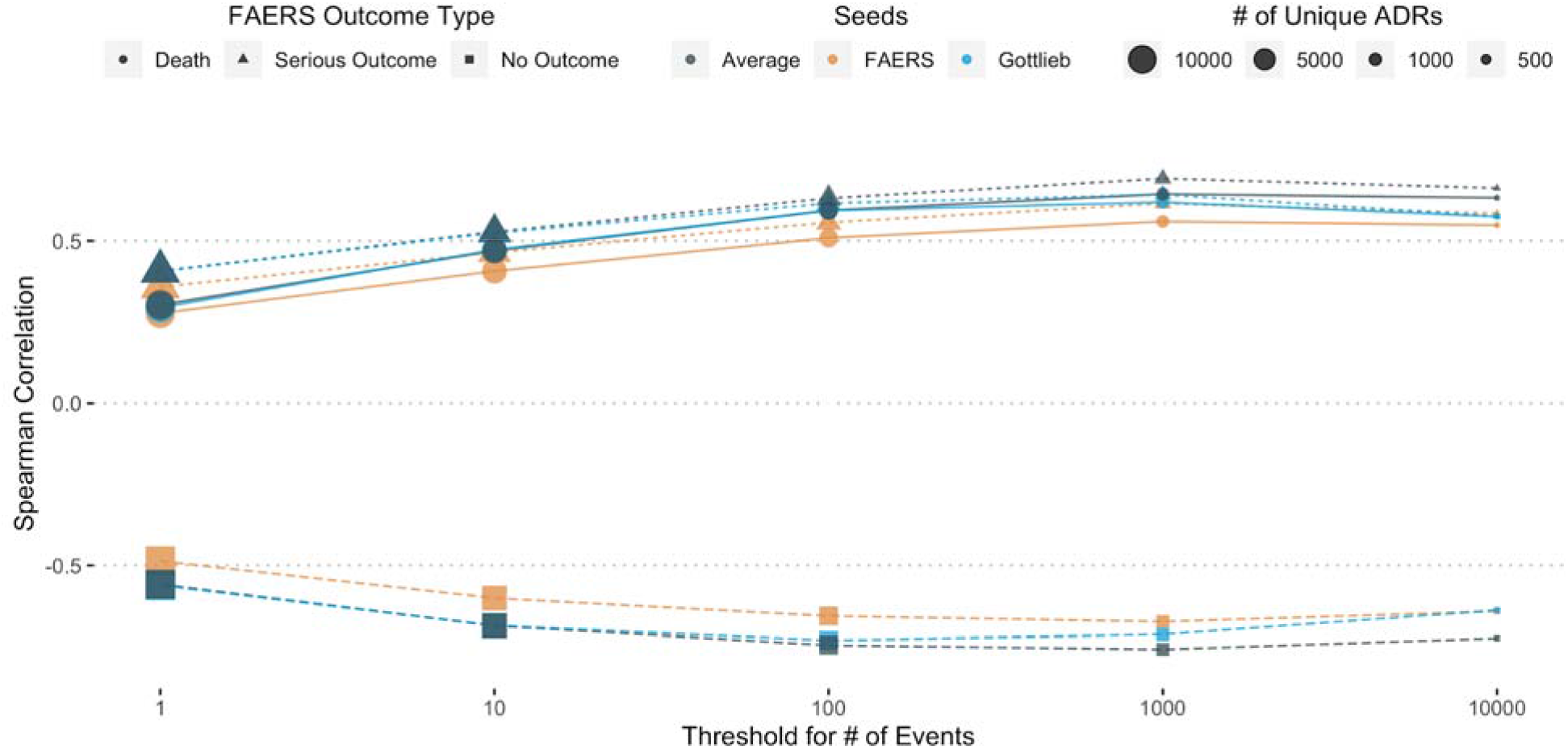
Spearman correlation of severity estimates with real-world outcomes in FAERS cases for differently case count thresholds. Comparing the performance of the different severity estimates across ADRs thresholded at various levels of number of events reported. Performance plateaus after >100 events for all estimates. The combined severity estimate has the best performance across all three outcome types.

**Supplementary Figure 2.**
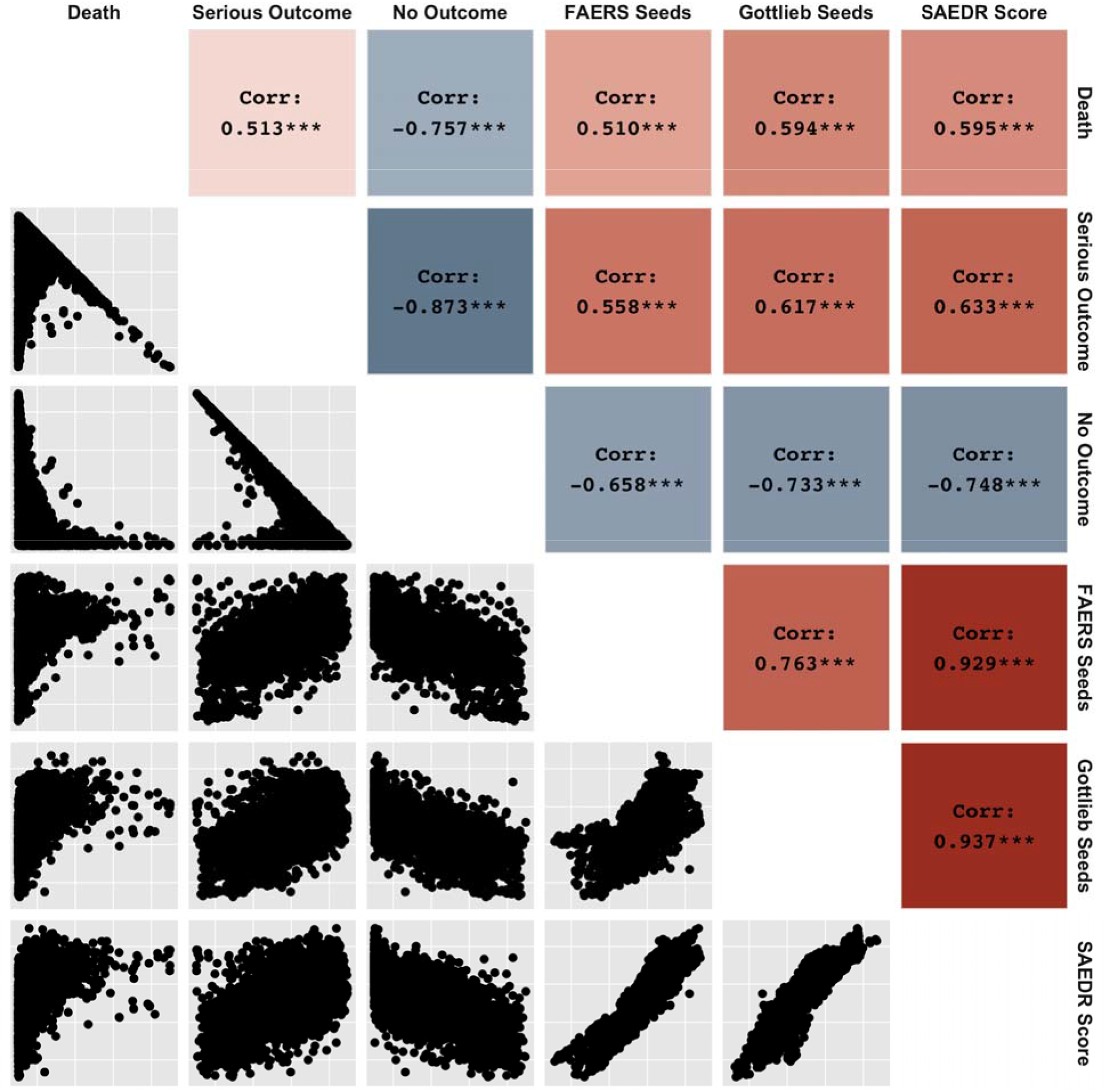
Pairwise plots and Spearman correlations between FAERS outcomes and different ADR severity estimates. Upper triangle shows pairwise Spearman correlations between FAERS outcomes of Death, Serious Outcome, and No Outcome and differently seeded severity estimates, as well as the final SAEDR score metric. Lower triangle shows pairwise plots for each set of scores, with each point representing a unique ADR.

**Supplementary Figure 3.**
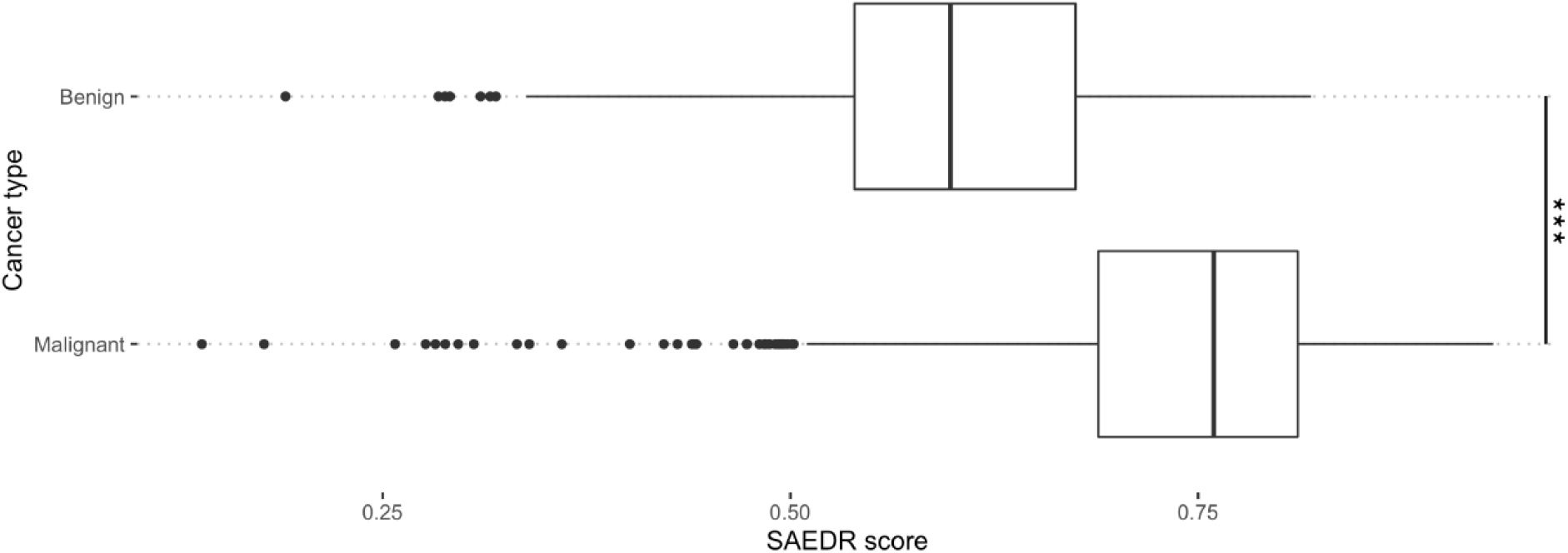
Differences in severity of neoplasm ADRs that contain the terms benign versus malignant. Malignant neoplasm ADRs had significantly higher severity scores than those associated with benign neoplasms, based on a one-sided t-test.

## References

1. Bourgeois FT, Shannon MW, Valim C. Adverse drug events in the outpatient setting: an 11 □year national analysis. and drug safety [Internet] Wiley Online Library; 2010; Available from: https://onlinelibrary.wiley.com/doi/abs/10.1002/

2. Watanabe JH, McInnis T, Hirsch JD. Cost of Prescription Drug-Related Morbidity and Mortality. Ann Pharmacother 2018 Sep;52(9):829–837. PMID:29577766

3. Aronson JK, Ferner RE. Clarification of terminology in drug safety. Drug Saf Springer; 2005;28(10):851–870. PMID:16180936

4. Ahmad SR. Adverse drug event monitoring at the Food and Drug Administration. J Gen Intern Med 2003 Jan;18(1):57–60. PMID:12534765

5. Code of Federal Regulations [Internet]. Code of Federal Regulations - Title 21 - Food and Drugs. Sect 21CFR20157. Available from: https://www.accessdata.fda.gov/scripts/cdrh/cfdocs/cfcfr/cfrsearch.cfm?fr=201.57

6. Shaffer ML, Watterberg KL. Joint distribution approaches to simultaneously quantifying benefit and risk. BMC Med Res Methodol 2006 Oct 12;6:48. PMID:17038184

7. Tallarida RJ, Murray RB, Eiben C. A scale for assessing the severity of diseases and adverse drug reactions. Application to drug benefit and risk. Clin Pharmacol Ther Wiley Online Library; 1979 Apr;25(4):381–390. PMID:428184

8. Gottlieb A, Hoehndorf R, Dumontier M, Altman RB. Ranking adverse drug reactions with crowdsourcing. J Med Internet Res 2015 Mar 23;17(3):e80. PMID:25800813

9. Mikolov T, Chen K, Corrado G, Dean J. Efficient Estimation of Word Representations in Vector Space [Internet]. arXiv [csCL]. 2013. Available from: http://arxiv.org/abs/1301.3781

10. Pennington J, Socher R, Manning CD. Glove: Global vectors for word representation. Proceedings of the 2014 conference on empirical methods in natural language processing (EMNLP) 2014. p. 1532–1543.

11. Firth JR. 1968. A synopsis of linguistic theory 1930-1955. Selected papers of JR Firth 1952-1959 1957;168–205.

12. Nikfarjam A, Sarker A, O’Connor K, Ginn R, Gonzalez G. Pharmacovigilance from social media: mining adverse drug reaction mentions using sequence labeling with word embedding cluster features. J Am Med Inform Assoc 2015 May;22(3):671–681. PMID:25755127

13. Hanson CL, Cannon B, Burton S, Giraud-Carrier C. An exploration of social circles and prescription drug abuse through Twitter. J Med Internet Res 2013 Sep 6;15(9):e189. PMID:24014109

14. Yan LL, Yan X, Tan Y, Sun SX. Shared minds: How patients use collaborative information sharing via social media platforms. Prod Oper Manag Wiley; 2019 Jan;28(1):9–26.

15. Alvaro N, Conway M, Doan S, Lofi C, Overington J, Collier N. Crowdsourcing Twitter annotations to identify first-hand experiences of prescription drug use. J Biomed Inform 2015 Dec;58:280–287. PMID:26556646

16. Klein A, Sarker A, Rouhizadeh M, O’Connor K, Gonzalez G. Detecting Personal Medication Intake in Twitter: An Annotated Corpus and Baseline Classification System. BioNLP 2017 Vancouver, Canada,: Association for Computational Linguistics; 2017. p. 136–142.

17. Lavertu A, Altman RB. RedMed: Extending drug lexicons for social media applications. J Biomed Inform 2019 Nov;99:103307. PMID:31627020

18. Kuhn M, Letunic I, Jensen LJ, Bork P. The SIDER database of drugs and side effects. Nucleic Acids Res 2016 Jan 4;44(D1):D1075–9. PMID:26481350

19. Brown EG, Wood L, Wood S. The Medical Dictionary for Regulatory Activities (MedDRA) [Internet]. Drug Safety. 1999. p. 109–117. [doi:10.2165/00002018-199920020-00002]

20. Bodenreider O. The Unified Medical Language System (UMLS): integrating biomedical terminology. Nucleic Acids Res 2004 Jan 1;32(Database issue):D267–70. PMID:14681409

21. Hamilton WL, Clark K, Leskovec J, Jurafsky D. Inducing Domain-Specific Sentiment Lexicons from Unlabeled Corpora. Proc Conf Empir Methods Nat Lang Process 2016 Nov;2016:595–605. PMID:28660257

22. Grover A, Leskovec J. node2vec: Scalable Feature Learning for Networks. KDD 2016 Aug;2016:855–864. PMID:27853626

23. Efron B. Bootstrap Methods: Another Look at the Jackknife. Ann Stat Institute of Mathematical Statistics; 1979 Jan;7(1):1–26.

24. Wu L, Ingle T, Liu Z, Zhao-Wong A, Harris S, Thakkar S, Zhou G, Yang J, Xu J, Mehta D, Ge W, Tong W, Fang H. Study of serious adverse drug reactions using FDA-approved drug labeling and MedDRA. BMC Bioinformatics 2019 Mar 14;20(Suppl 2):97. PMID:30871458

25. Chandak P, Tatonetti NP. Using Machine Learning to Identify Adverse Drug Effects Posing Increased Risk to Women. Patterns (N Y) [Internet] 2020 Oct 9;1(7). PMID:33179017

26. Tatonetti NP, Ye PP, Daneshjou R, Altman RB. Data-driven prediction of drug effects and interactions. Sci Transl Med 2012 Mar 14;4(125):125ra31. PMID:22422992

27. Watson S, Caster O, Rochon PA, den Ruijter H. Reported adverse drug reactions in women and men: Aggregated evidence from globally collected individual case reports during half a century. EClinicalMedicine 2019 Dec;17:100188. PMID:31891132

28. Holm L, Ekman E, Jorsäter Blomgren K. Influence of age, sex and seriousness on reporting of adverse drug reactions in Sweden. Pharmacoepidemiol Drug Saf 2017 Mar;26(3):335–343. PMID:28071845

29. Lavertu A, Vora B, Giacomini KM, Altman R, Rensi S. A New Era in Pharmacovigilance: Towards real world data and digital monitoring. Clin Pharmacol Ther [Internet] Wiley; 2021 Jan 25;(cpt.2172). [doi:10.1002/cpt.2172]

30. Pinnow E, Amr S, Bentzen SM, Brajovic S, Hungerford L, St George DM, Dal Pan G. Postmarket Safety Outcomes for New Molecular Entity (NME) Drugs Approved by the Food and Drug Administration Between 2002 and 2014. Clin Pharmacol Ther 2018 Aug;104(2):390–400. PMID:29266187

31. Quinn KJ, Shah NH. A dataset quantifying polypharmacy in the United States. Sci Data 2017 Oct 31;4:170167. PMID:29087369

32. Hazell L, Shakir SAW. Under-reporting of adverse drug reactions: a systematic review. Drug Saf Springer; 2006;29(5):385–396. PMID:16689555

